# Auditory sensory memory span for duration is severely curtailed in females with Rett Syndrome

**DOI:** 10.1101/568832

**Authors:** Tufikameni Brima, Sophie Molholm, Ciara J. Molloy, Olga V. Sysoeva, Eric Nicholas, Aleksandra Djukic, Edward G. Freedman, John J. Foxe

**Author notes:** Corresponding Author: John J. Foxe, Ph.D., Department of Neuroscience, University of Rochester Medical Center, 601 Elmwood Avenue - Box 603, Rochester, New York 14642.

## Abstract

Rett syndrome (RTT), a rare neurodevelopmental disorder caused by mutations in the *MECP2* gene, is typified by profound cognitive impairment and severe language impairment, rendering it very difficult to accurately measure auditory processing capabilities behaviorally in this population. Here we leverage the mismatch negativity (MMN) component of the event-related potential to measure the ability of RTT patients to decode and store occasional duration deviations in a stream of auditory stimuli. Sensory memory for duration, crucial for speech comprehension, has not been studied in RTT.

High-density EEG was successfully recorded in 18 females with RTT and 27 age-matched typically developing (TD) controls (aged 6-22 years). Data from 7 RTT and 3 TD participants were excluded for excessive noise. Stimuli were 1kHz tones with a standard duration of 100ms and deviant duration of 180ms. To assess the sustainability of sensory memory, stimulus presentation rate was varied with stimulus onset asynchronies (SOAs) of 450, 900 and 1800ms. MMNs with maximum negativity over fronto-central scalp and a latency of 220-230ms were clearly evident for each presentation rate in the TD group, but only for the shortest SOA in the RTT group. Repeated-measures ANOVA revealed a significant group by SOA interaction. MMN amplitude correlated with age in the TD group only. MMN amplitude was not correlated with the Rett Syndrome Severity Scale. This study indicates that while RTT patients can decode deviations in auditory duration, the span of this sensory memory system is severely foreshortened, with likely implications for speech decoding abilities.

## INTRODUCTION

Rett syndrome (RTT) is a severely incapacitating neurodevelopmental disorder affecting about 1-in-10,000 females [1–3]. RTT typically results from spontaneous mutations in the X-linked gene encoding Methyl-CpG-binding protein 2 (*MeCP2*) [2]. Clinically, RTT is characterized as an age-specific (6-18 months) progressive loss of initially acquired intellectual, language and motor abilities, with loss of purposeful hand use that is often supplanted by distinctive hand stereotypies and gait abnormalities [1, 3]. Very little is known about cognitive processing, including perceptual capabilities and speech comprehension across the progressive clinical stages of RTT [4, 5]. This is largely due to the nature of the impairments associated with RTT, which typically preclude the use of conventional cognitive evaluations which require verbal and/or gestural responses [6]. This leaves the field with non-representative outcome measures, making it difficult to assess neurocognitive function during the natural course of RTT or in response to treatments during clinical trials [7]. As such, there is an urgent need to develop objective quantitative measures of brain function (i.e. neural markers) that can be tracked in a noninvasive and unbiased manner.

Event related potential (ERP) recordings are an increasingly appealing option in both patients with RTT and animal models as a means to assess information processing and cognitive capabilities [8–14]. This technique provides the opportunity to deliver objective quantitative measures of brain function, including cortical network dynamics, in the absence of overt behavioral responses from participants (e.g. [15–17]). The integrity of early auditory processing, auditory discrimination and sensory memory can be studied with the ERP component known as mismatch negativity (MMN) [18, 19]. MMN is a relatively automatic ERP response evoked by occasional changes in a regularly occurring stream of inputs. Assessing MMN is ideal for RTT in that no engagement from the listener is required.

In a recent study, we used high-density ERPs to examine the ability of individuals with RTT to automatically process pitch changes in a regularly occurring stream of auditory tones [9]. Accurate neural representation of time-varying spectral changes in the auditory system is clearly a critical component of processing the speech signal [20]. Individuals with RTT showed severely impaired basic auditory sensory processing such that their automatic representation of pitch discrimination in sensory memory was delayed and prolonged relative to their typically developing peers. However, the presence of a MMN response, although morphologically atypical and delayed, suggested that these individuals are capable of detecting frequency deviations within a continuous stream of auditory inputs.

To build on these previous findings, the current study sought to further characterize auditory sensory memory capabilities in RTT, by specifically exploring whether there is adequate cortical representation of auditory duration, another critical cue in speech perception [21]. This is the first study to examine this important characteristic of sensory memory in RTT. To further interrogate the robustness of sensory memory for duration, we examined whether these sensory memory traces could be sustained over longer and longer periods. It has been shown that the robustness of the memory trace represented by the MMN is directly related to the rate at which the regularly occurring stream of stimuli is presented [22, 23]. To test this in RTT, we assessed the MMN for duration deviance using three stimulation rates. We hypothesized that MMN generation would be present in RTT individuals, but that it would be more severely impacted over slower stimulation rates.

## METHODS

### Participants

Twenty-five patients with Rett syndrome (RTT), confirmed by *MECP2* mutations, and thirty age-matched neurologically typically developing (TD) individuals took part in the study. Participants with RTT were recruited through the Rett Syndrome Center of the Children’s Hospital at Montefiore, while TD participants were recruited from the local community. All RTT patients were female as this rare disease affects mostly females. Ten out of thirty TD participants were male. Seven data sets from RTT and three from the TD group were excluded from analysis due to noisy EEG data resulting in less than 20% accepted trials per condition. The final sample contained eighteen females with RTT (mean age: 12.8±4.7, ranged 6-22) and twenty-seven TD controls (12.1±4.8, ranged 6-26). There was no significant difference in age between the RTT and TD group (t(43) = 0.5, p = 0.61). All participants with RTT underwent genetic testing, and detailed phenotypic assessment that was accompanied by detailed medical history questionnaires completed by their caregivers. Symptom severity in RTT was measured using the Rett Syndrome Severity Scale (RSSS) [24]. This clinician-rated scale represents an aggregate measure of the severity of clinical symptoms, including motor function, seizures, respiratory irregularities, ambulation, scoliosis, and speech. Composite scores in the range of 0 – 7 correspond to a mild phenotype, 8 - 14 correspond to a moderate phenotype, and 15 - 21 to severe features. The RSSS score in the current RTT group ranged between 5 and 15 (Mean ± SD = 9.6±4). Clinical demographics such as severity scores, ages (age of onset and regression) of all participants including those with unusable data and medication are listed in supplementary material (Supplementary Table 1 C*linical Demographics*). There were no differences in age-range and RSSS scores between the 7 excluded RTT datasets and those included in the final analysis.

Participants with RTT were excluded if they were experiencing ongoing regression, specifically in Stage II, which is also known as the rapid developmental regression stage. Other exclusion criteria included uncorrected hearing loss or ear infection on the day of EEG acquisition. TD participants were excluded if they had a familial history of a neurodevelopmental disorder, any neurological or psychiatric disorders. All individuals in the TD group passed a hearing screen. While none of the RTT participants were deaf, hearing acuity could not be similarly assessed in the RTT participants.

The institutional review boards of the University of Rochester and Albert Einstein College of Medicine approved this study. Written informed consent was obtained from parents or legal guardians. Where possible, informed assent from the participants was also obtained. Participants were modestly compensated at a rate of $15 per hour for their time in the laboratory. All aspects of the research conformed to the tenets of the Declaration of Helsinki.

### Experimental Design

We presented a simple auditory MMN paradigm while recording high-density electroencephalography (EEG) from TD and RTT participants. Experimental procedures were similar to those described in our previous paper [9]. All participants sat in a sound-attenuated and electrically-shielded booth (Industrial Acoustics Company, Bronx, NY) on a caregiver’s lap or in a chair/wheelchair. They watched a muted movie of their choice on a laptop (Dell Latitude E640) while passively listening to auditory stimuli presented at an intensity of 75 dB SPL using a pair of Etymotic insert earphones (Etymotic Research, Inc., Elk Grove Village, IL, USA). An oddball paradigm was implemented in which regularly occurring standard tones (85%) were randomly interspersed with deviant tones (15%). These tones had a frequency of 1000 Hz with a rise and fall time of 10 ms. Standard tones had duration of 100 ms while deviant tones were 180 ms in duration. The tones were presented with stimulus onset asynchronies (SOAs) of either 450, 900 or 1800 ms in separate blocks, and each block consisted of 500, 250 or 125 trials respectively (supplementary Fig 1A). Participants were presented with a total of 14 blocks (2×450ms, 4×900ms and 8×1800ms) equally distributed within the experimental session, resulting in 1000 trials per condition.

**Figure 1.**
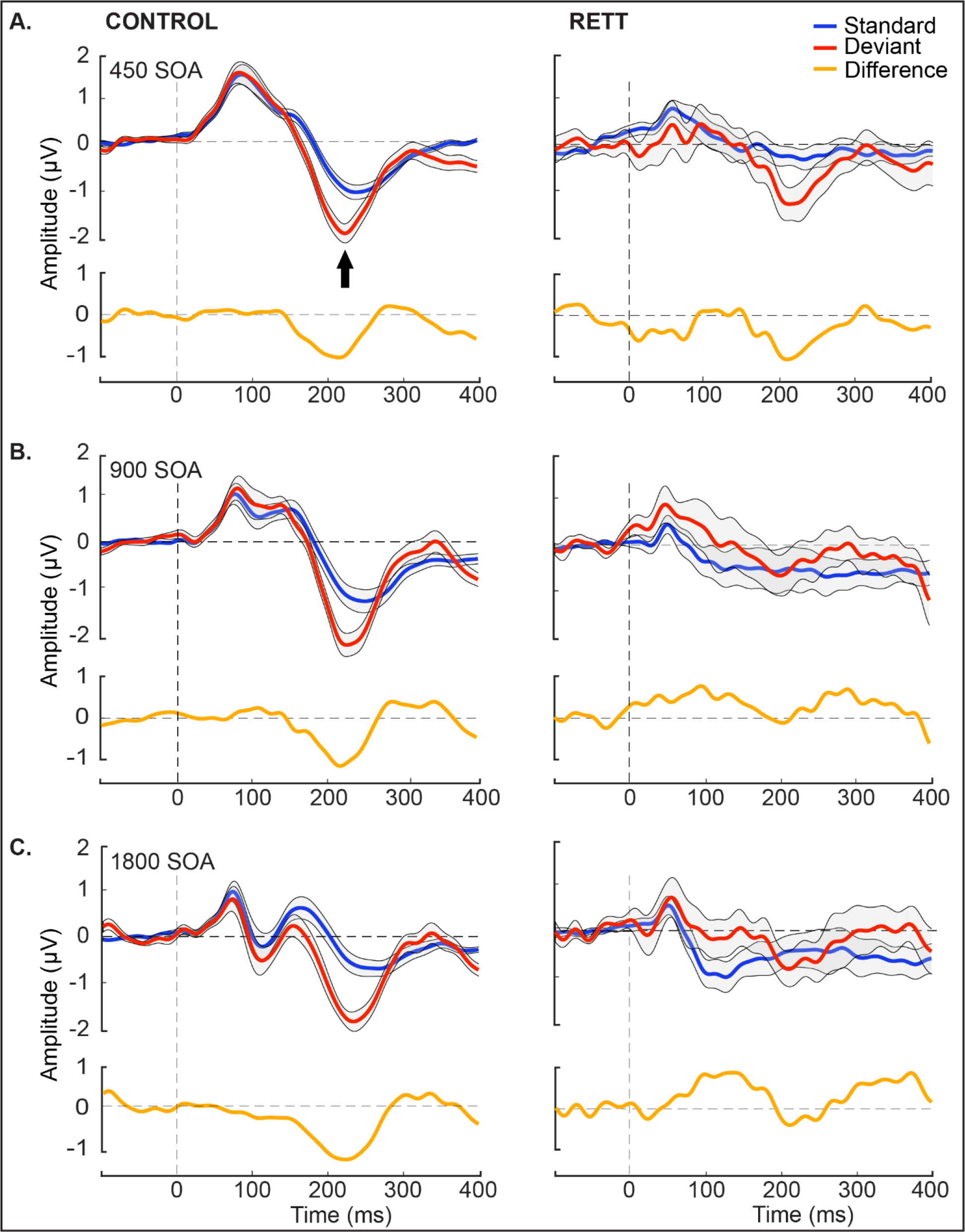
Grand mean waveforms for TD and RTT group over fronto-central electrodes (FC3, FCz and FC4). Auditory event-related potentials (ERPs) to standard tones (blue trace) and deviant tones (red trace) are presented with standard deviation, shaded in gray. The TD group produced classic ERP waveforms, while the RTT group exhibited less stereotyped responses with reduced ERP amplitude across conditions. A clear MMN (difference between standard and deviant traces) was present for all SOAs for the TD group. However, an MMN was present only for the shortest SOA in the RTT group.

### EEG Acquisition

A Biosemi ActiveTwo (Bio Semi B.V., Amsterdam, Netherlands) 72-electrode array was used to record continuous EEG signals. The set up includes an analog-to digital converter, and fiber-optic pass-through to a dedicated acquisition computer (digitized at 512 Hz; DC-to-150 Hz pass-band). EEG data were referenced to an active common mode sense (CMS) electrode and a passive driven right leg (DRL) electrode.

### EEG Data processing

EEG data were processed and analyzed offline using custom scripts that included functions from the EEGLAB Toolbox [25] for MATLAB (the MathWorks, Natick, MA, USA). EEG data were initially high-pass filtered using a Chebyshev Type II filter with a bandpass set at 1-40 Hz. Continuous EEG data were passed through a channel rejection algorithm, which identified bad channels using measures of standard deviation and covariance with neighboring channels. Rejected channels were interpolated using the EEGLAB spherical interpolation. Data were then divided into epochs that started 100 ms before the presentation of each tone and extended to 800 ms post stimulus onset. Bad trials containing severe movement artifacts or particularly noisy events were rejected if voltages exceeding ±150 μV, followed by a threshold set at two standard deviations over the mean of the maximum values for each epoch (the largest absolute value recorded in the first 500 ms of a given epoch, across all channels for each trial in each condition). The number of accepted trails for each condition and group is presented in supplementary figure 2. All epochs were then baseline corrected to the 100 ms pre-stimulus interval. The epochs were next averaged as a function of stimulus condition to yield the auditory evoked potential to the standard and to the deviant tone. To maximize the ERP at fronto-central sites, the data were referenced to TP7, or TP8 if TP7 was a noisy channel in a given participant. This approach takes advantage of the inversion of the MMN that is seen between fronto-central and inferior temporo-parietal sites [26–28].

**Figure 2.**
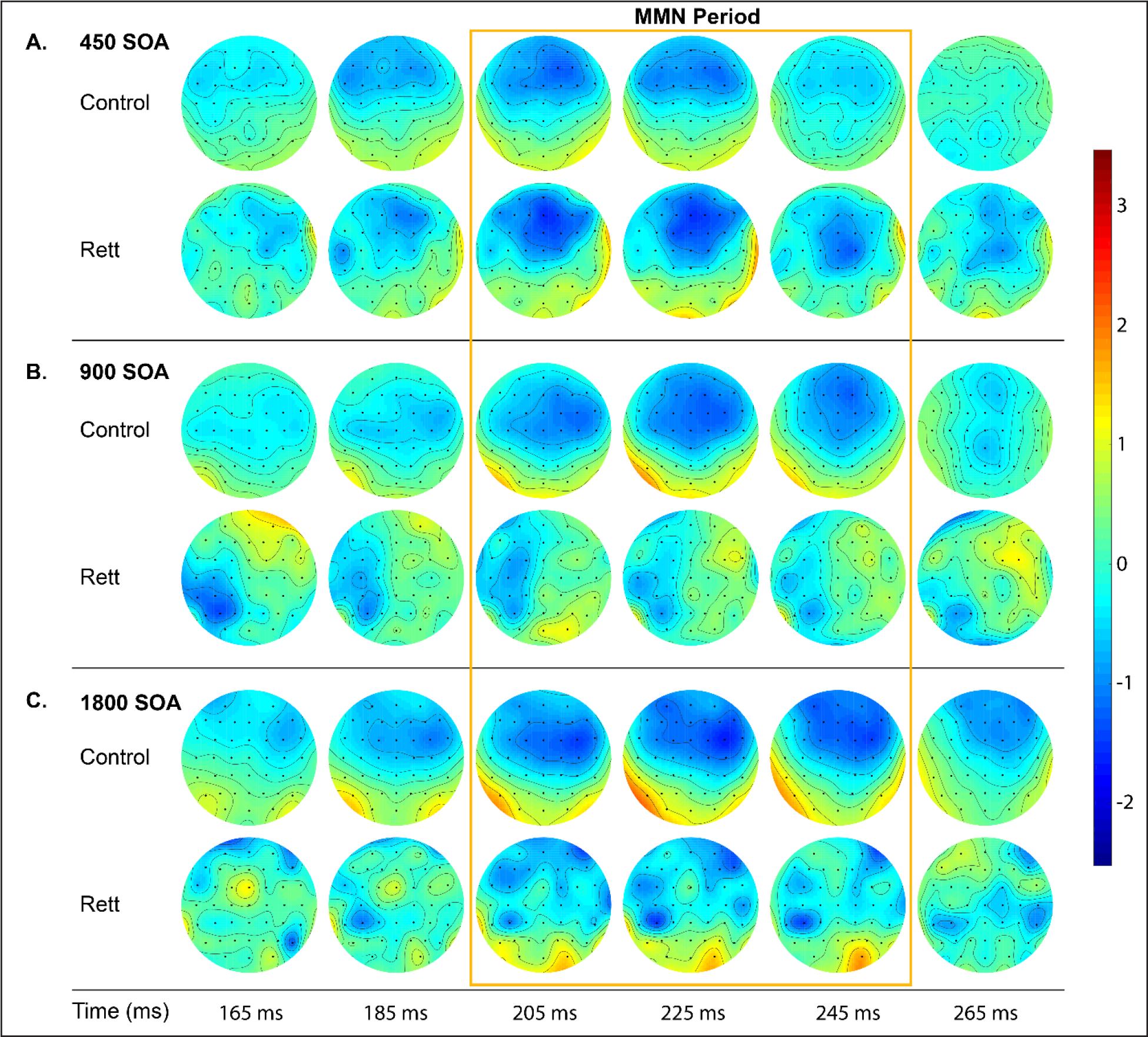
Topographic representation of the differences between deviant and standard tones across SOAs. An MMN with typical spatial distribution with negativity at the frontal site and positivity over the mastoids is clearly seen in all conditions for TD group, but only for the shortest SOA condition in the RTT group.

The window for measurement of the MMN was calculated by subtracting the grand mean ERP to deviant tones from the grand mean ERP to standard tones. The resulting distribution of activity showed maximal difference at approximately 225 ms (Fig 2 A-C). We then defined a time window of 10 ms centered around 225 ms (i.e., 220-230 ms) to obtain average MMN amplitudes for each individual for each SOA. Composite averages generated from FC3, FCz and FC4 scalp electrodes were used for further statistical analysis.

### Statistical analyses

The primary analysis employed a repeated measures analysis of variance (ANOVA) with SOA (450, 900 and 1800 ms) as a within-participant factor and Group (RTT vs. TD) as a between-participants factor to examine main effects and their interaction on MMN amplitude. Planned *post-hoc* tests were used to follow significant ANOVA effects. Partial η^2^ was used to estimate effect sizes. Pearson correlations were used to assess the relationship between MMN amplitude, age and RSSS. We also compared the correlation coefficients of age with the MMN for each SOA using a Fisher z-transformation to examine if MMN maturation was similar across SOAs [29].

In a secondary exploratory analysis, we further interrogated this rich high-density EEG dataset using the statistical cluster plot (SCP) approach (see Figure 4). This approach serves as a follow-up to the primary a-priori tests of the MMN, as a means to more fully describe the recorded data, and as such, it is to be considered exploratory. Any additional effects uncovered should be considered post-hoc, must be interpreted cautiously, and serve simply as hypothesis generation tools to be explored further in a follow-up study. SCPs are constructed by calculating pairwise 2-tailed t-tests between the evoked responses to a given pair of experimental conditions across all timepoints and all recording sites (electrodes). Results of these tests are then displayed as a color intensity plot that spatio-temporally summarizes periods of statistical difference between conditions (here: standard versus deviant). The X and Y axes signify time (in milliseconds) and electrode location (frontal to posterior scalp) respectively, while the color represents the t-value for each data point. Only points exceeding an alpha criterion of .05 or less are highlighted, and then only when this criterion is present for a minimum of 11 consecutive data points (i.e. 21.5 ms at the current digitization rate), thereby reducing the probability of type I errors. That is, the likelihood of multiple false positive results occurring by chance at 11 consecutive time points is exceedingly low if one assumes statistical independence between each time point. Of course, there is a degree of autocorrelation between temporally adjacent time points in EEG recordings that must be considered. Even for high autocorrelations, applying a criterion of 11 consecutive time points has been shown to be quite conservative in avoiding type I errors [30, 31]

## RESULTS

Figure 1 displays ERPs elicited by standard and deviant tones for each SOA, as well as the corresponding difference waves, over fronto-central scalp sites (the average of FC3, FCz and FC4). Clear MMN responses can be seen for every SOA in TD controls, while this is only the case for the shortest SOA (450ms) in the RTT group. These MMNs have typical topography with negativity over the fronto-central sites and positivity in the mastoids as can be seen on Figure 2. MMNs lasted from approximately 180 to 250 ms with maximum at about 225 ms, so the average MMN amplitude within 220-230 ms latencies was chosen for statistical analyses. Repeated measures ANOVA revealed a main effect of Group (F(1,43) = 6.970, *p* = 0.012, η^2^ = 0.139), indicating the attenuated MMN in RTT as compared to TD. The main effect of SOA did not reach significance (F(2,42) = 1.897, *p* = 0.156, η^2^ interaction did (F(2,42) = 4.348, *p* = 0.017, η^2^ = 0.042), whereas the Group by SOA = 0.092, Greenhouse-Geisser correction applied) confirming the evident reduction of MMN in RTT at slower presentation rates (i.e. SOAs of 900 and 1800 ms). *Post-hoc* analyses also confirmed that TD and RTT groups differed significantly only for the longer SOAs of 900 ms (F(1,43) = 6.728, p = 0.013, η^2^ = 0.135) and 1800 ms (F(1,43) = 10.302, p = 0.003, η^2^ = 0.193), but not for the shortest SOA of 450 ms (F(1,43) = 0.046, p = 0.831, η^2^ = 0.001). The mean MMN amplitude for the different SOAs in each group can be seen in Figure 3.

**Figure 3.**
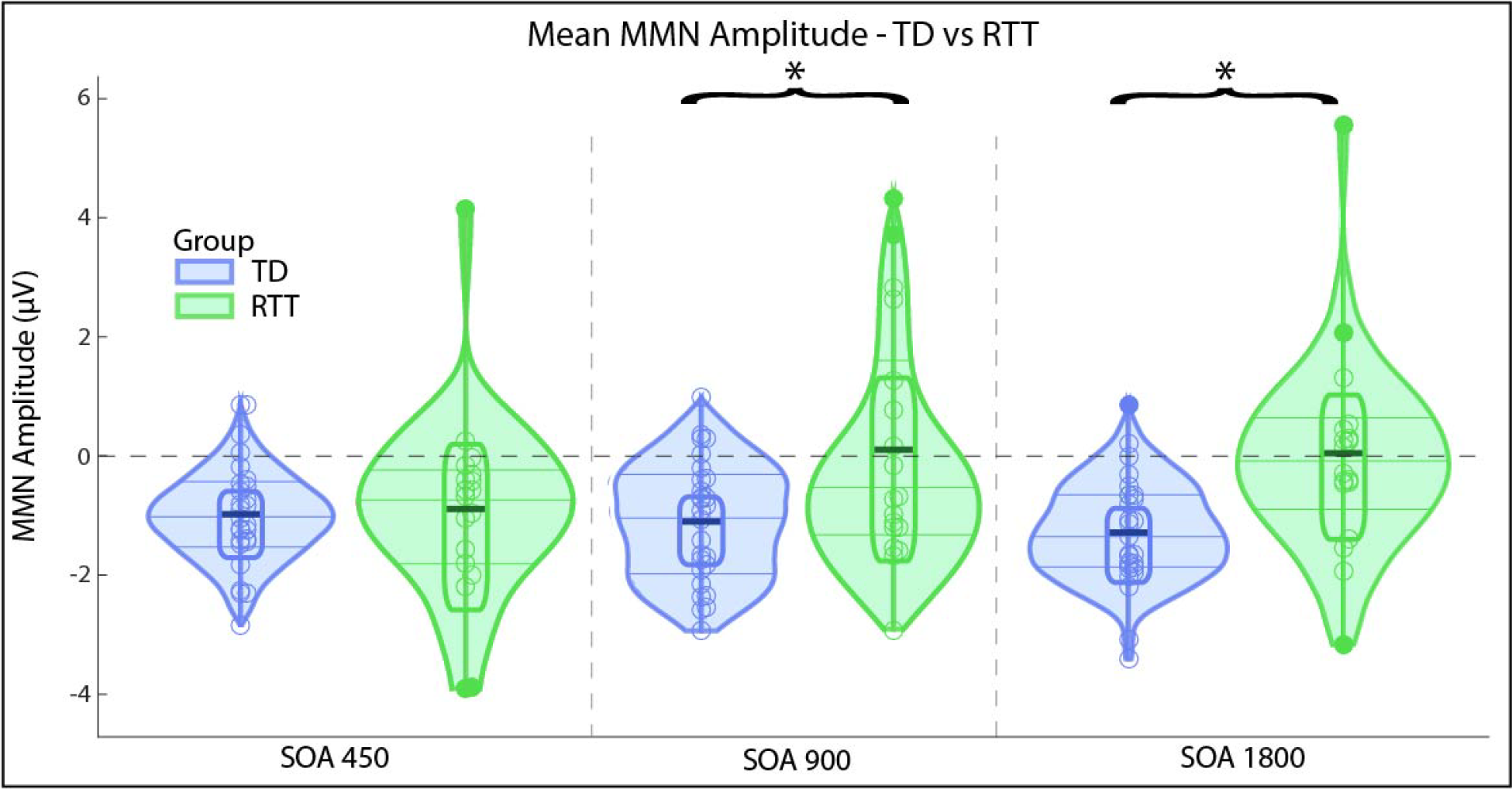
Mean MMN amplitude for each SOA in RTT and TD groups. Significant difference between the groups are marked by asterisk (SOAs of 900 and 1800 ms).

**Figure 4.**
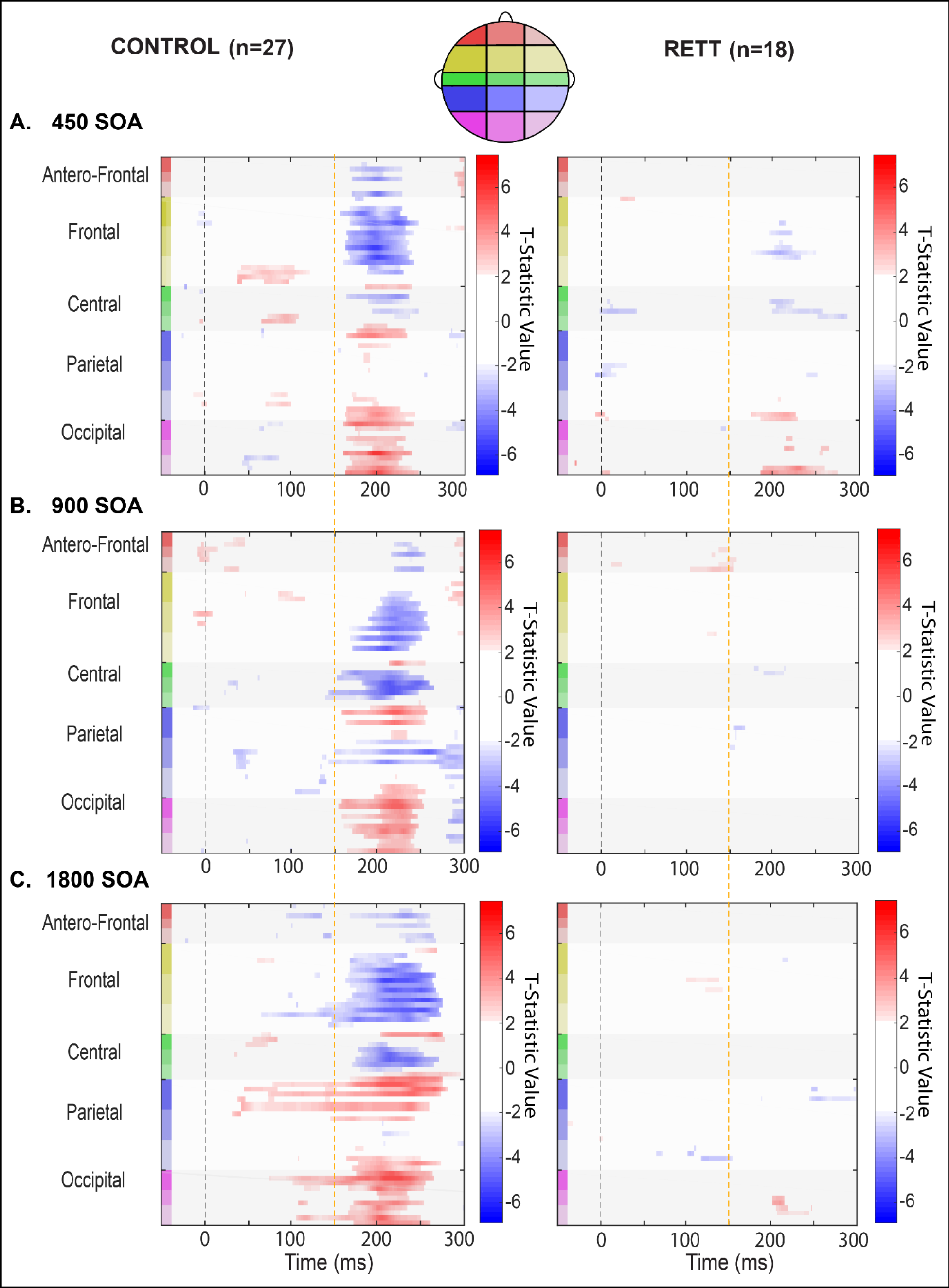
Statistical cluster plots (SCPs) depicting the outcome of running t-tests that compare standard and deviant responses generated by each group (TD and RETT) to each condition (450 ms, 900 ms and 1800 ms SOAs), for all electrodes and all time points (−50 ms to 300 ms). Significant effects are plotted when p ≤ 0.05 for at least 9 consecutive data points (~18 ms at 512 Hz sampling rate). The direction of these effects is color coded: When responses to deviant tones are significantly more positive (red) or significantly negative (blue) relative to responses to standard tones. The grey areas indicate periods where no differences are observed. Time is plotted along the x–axis and electrode position is plotted on the y-axis. Starting from the top left corner of each graph, electrodes that are located next to each other are clustered into color-coded scalp regions (Frontal, Central, Temporal, and Occipital). These color-coded regions are displayed on the corresponding head map (right).

MMN amplitude showed a significant negative linear correlation with age in the TD group, for SOAs of 450 and 900 ms (r(27) = −0.41, p(27) = 0.035 and r(27) = −0.39, p(27) = 0.045, respectively), and at a trend level for SOA of 1800 ms (r(27) = −0.32, p(27) = 0.1). There were no differences in correlation coefficients across SOAs in TD (z = −0.4295, p-value = 0.6675). For the RTT group, MMN amplitude was not linearly correlated with age (|r| < 0.21, p(18) > 0.4) or RSSS (|r| < 0.21, p(18) > 0.4).

In follow-up exploratory analysis, the SCP approach was used to assess the general time-course of significant MMN differences between conditions (i.e. standard versus deviant) for each of the three presentation rates and for both groups (Figure 4). As can be seen in the left panels of the figure, a robust differential effect was seen at all three presentation rates for TD participants, confirming and extending the findings of the ANOVA reported above. In contrast, only at the fastest presentation rate (450 ms) was a differential effect evident in the RTT population, with a considerably later onset (~190 ms versus 160 ms) and somewhat shorter-lived duration of differential significance over a much more circumscribed set of channels. No effects of note were evident at the two slower presentation rates in the RTT group, again confirming the main findings of the ANOVA above.

## DISCUSSION

The current data show that participants with RTT produce the MMN response for the shortest 450 ms SOA condition, albeit attenuated and delayed, indicating that they have somewhat preserved ability to build an auditory sensory memory representation for duration and to discriminate duration deviance when a sequence of auditory signals is presented at a fast stimulation rate. However, these sensory memory traces for tone duration appear to have a substantially foreshortened temporal span in RTT, as evident by the highly atypical absence of a detectable MMN at longer SOAs of 900 and 1800 ms.

The presence of an MMN to auditory duration deviance in the current study corresponds with our previous findings using a pitch deviance paradigm, where the presence of an MMN pointed to the neural ability to represent frequency deviations in RTT albeit that the pitch-evoked MMN was morphologically atypical and substantially delayed in that study [9]. The presence of a duration-evoked MMN here similarly suggests the neural capacity to present duration deviants. On the other hand, that the duration MMN is absent at slower presentation rates, might also suggest that sensory memory for this fundamental auditory feature is more impacted in RTT than is frequency representation. It is worth noting, however, that in our prior study, only a single frequency manipulation was used to generate the MMN, and that the difference between the standard and deviant (503 versus 996Hz) was very large and therefore highly detectable. Unlike the current study, we did not parametrically vary the strength of the memory trace, either by changing the presentation rate or by narrowing the frequency difference. Taken together, the two studies suggest that while MMN is relatively preserved in RTT when the stimulation parameters are such that the deviation, be it in duration or pitch, is large and highly detectable, once the system is taxed to any degree, significant deficits begin to reveal themselves. Clearly, it will be important to follow up with parametric studies to assess the limits of the auditory sensory memory system in RTT for these and other fundamental auditory features (i.e., frequency, duration, location and loudness).

It is also worth pointing out that the specificity of impaired MMN to the longer SOA conditions indicates that the MMN reduction was unlikely caused by factors such as attention or motivation, which under certain limited circumstances are known to modulate MMN amplitude [32]. The intact MMN in the fastest condition, and the fact that the blocks with different SOAs were equally distributed within experimental session clearly show that it is the ability to sustain sensory memory representations of duration over longer periods that is reduced in RTT.

Our study also showed an increase in MMN amplitude with age in the TD group, consistent with previous findings [33]. Noteworthy, the rate of MMN maturation was similar across SOAs, indicating that while duration discrimination ability is still developing from childhood into adulthood, the ability to sustain auditory sensory memory representations is already above 2 seconds within the age range of the participants in this study (6-26 years old). This result is in line with prior work showing that auditory sensory memory for frequency can last for at least 3 seconds in 6 year old children [34]. In typical development, there is a gradual increase in the duration of sensory memory with development, such that no MMN is observed in children younger than 2 years old when SOAs are longer than 1 second, in 4-year-old children when SOAs are longer than 2 seconds, or in 6-year-old children with SOAs longer than 3 seconds. However, the MMN is prominent in each of these groups with faster presentation rates [34–36].

One possible implication of the fact that patients with RTT showed no reliable MMN at 900 ms, is that their sensory memory systems may have stagnated at a developmental stage compatible with that of a 1-2-year-old TD child. It is in this age range that patients with RTT usually experience developmental arrest that affects cognitive and motor functions, including language skills. Another measure of basic auditory function previously examined in RTT is the brainstem frequency following response, which measures responses that are phase-locked to the frequency of a periodic stimulus. Galbraith and colleagues, working with adults with RTT, found that the ability to entrain external high-frequency modulation was at a level more consistent with infant maturation levels seen in TDs [37]. Thus, it might be suggested that auditory system maturation is also halted at an early developmental stage.

One might argue that the decay in sensory memory could be specific to duration representation, since previous studies have suggested that memory representation for duration decays more rapidly with aging and cognitive decline than does frequency representation [38–40]. In our previous study of RTT patients, MMN data suggested that the lifetime of frequency representation is at least 900 ms, longer than observed here for duration, but as mentioned, that was the only SOA employed in that study [9]. Nonetheless, it is a distinct possibility that the duration system is more vulnerable than is the frequency representation system, and this may relate to the separable cortical circuits responsible for generating these two varieties of the MMN [26]. Whereas the major generators of the frequency MMN are found in hierarchically early primary auditory cortex as well as fronto-parietal regions, a more complex set of regions was involved in generating the duration MMN, including secondary auditory cortex and a more extended set of fronto-parietal generators. It is plausible that higher-order cortical regions are more impacted by the disease than are early regions, and that it follows that processes reliant on these higher-order regions might be more vulnerable. However, that we have not yet parametrically varied the presentation rates for frequency deviations within the same cohort, leaves this an open question for now.

The MMN measures in our study were not related to RTT severity as measured by the RSSS. While the RSSS accumulates all aspects of RTT symptomatology and is based on observation and clinical symptoms, the MMN is an objective measure of auditory sensory memory. The absence of correlation between the two measures suggests that cognitive and perceptual deficits, which are difficult to assess via behavioral methods, may be independent from core motor symptoms. Thus, our neurophysiological measure provides critical additional information about auditory perceptual and cognitive brain function in patients with RTT that is not directly related to the currently used clinical measures of RTT severity. Importantly, it also has implications for speech and language comprehension.

The attenuated memory span, as measured by MMN to stimulation with different SOAs is characteristic of several neurological conditions. MMN generation is crucially dependent on the proper functioning of NMDA receptors, as shown by animal studies as well as pharmacological studies in healthy adults and patients with schizophrenia [41, 42]. NMDA abnormalities have also been implicated in RTT. In a small post-mortem study that included 9 independent RTT samples from prefrontal cortex, Blue and colleagues (1999) reported that NMDA receptor density was increased in samples from younger girls (2-8 years old), while it was decreased in samples from older girls (up to 30 years of age) [43]. Similar biphasic shifts in the direction of this NMDA effect with age were confirmed in prefrontal brain regions of a RTT animal model [44]. While there was no age-related difference in MMN changes across condition in our RTT patients, most of them were older than 8 years. NMDA receptor dysfunction has also been reported in healthy elderly subjects and those with Alzheimer’s disease [45], which have both also been shown to exhibit shortened sensory memory [39, 46–48]. Therefore, the absence of MMN in RTT for longer SOAs could, in theory, be linked to decreased NMDA receptor binding. However, there is no evidence that NMDA receptor dysfunction would affect MMN only in the two slower presentation rates, as patients with schizophrenia show reduced MMN even with short SOAs of 450-500 ms [49, 50]. Thus, understanding the role of NMDA receptors in modulating the span of sensory memory traces in different neurological disorders needs further investigation.

The oddball paradigm with varied SOAs can be easily implemented in animals. The animal model of RTT, either with total *Mecp2* knockout or deficient *Mecp2* in specific neuronal subtypes [11, 51], is a potentially very valuable tool to track down the neurophysiological and molecular mechanisms underlying the drastic decay in auditory sensory memory representation over longer periods that we report in this study, as well as to investigate its response to treatment. Taking into account that such profound deficits in sensory memory might be of crucial importance for language acquisition, the index of MMN attenuation with increasing SOA can be a very promising biomarker not only for RTT subjects, but for other neurodevelopmental disorders.

### Study Limitations

A limitation of the current study is the wide age-range of the participants, given that auditory responses continue to mature with typical development across the age-range tested [33, 52]. Although age was correlated with MMN amplitude in the TD group, it was not associated with manipulations of stimulus rate. This suggests that group differences seen as a function of presentation rate were not affected by age, but rather, represent frank differences in brain function in RTT. A second limitation is that mutation subtype could not be effectively examined here due to the relatively restricted sample size. Neither were we able to consider potential differences as a function of classic versus atypical Rett phenotype. Both of these distinctions will be of great interest as this work progresses. We must also acknowledge that hearing testing was not conducted at the time of EEG acquisition in patients with RTT due to difficulties in assessing it in this population. However, the presence of an MMN for the shortest SOA in the RTT group, clearly indicates that they could detect and decode auditory information. Lastly, non-invasive recordings such as those conducted here cannot shed light on the mechanisms by which MECP2 protein loss leads to auditory cortical processing deficits. Work using similar paradigms in murine models of RTT will be highly instructive in this regard [11, 53].

### Conclusions

This study confirms the preserved ability of RTT patients to automatically decode duration deviations in the auditory stream when stimuli are presented in a rapid stream, which we previously showed in this population for large frequency deviations. However, automatic detection of duration changes was highly atypical in RTT when the presentation rate of the stimulus stream was slowed down and the auditory sensory memory system was taxed, indicated by the lack of obvious MMN responses at SOAs of 900 and 1800 ms. We speculate that this drastic attenuation in the duration of sensory memory might lead to significant problems in language acquisition in RTT, as well as having implications for other aspects of information processing. The exact mechanisms underlying this decay, as well as behavioral outcomes, represent important avenues of research to increase knowledge of RTT and its perceptual and cognitive sequelae.

## Acknowledgments

This work was supported by a grant from the U.S. National Institute on Deafness and Other Communication Disorders (NIDCD R21 DC012447 to JJF and AD) and from The Rett Syndrome Research Trust. The authors acknowledge the participation of Mr. Douwe Horsthuis and Mr. Emmett Foxe during the data collection phase of this project. We also thank the volunteers and their families, who graciously gave of their time to allow for the successful completion of this study.

## Uncategorized References

1. Neul, J.L., et al., Rett syndrome: revised diagnostic criteria and nomenclature. Ann Neurol, 2010. 68(6): p. 944–50.

2. Amir, R.E., et al., Rett syndrome is caused by mutations in X-linked MECP2, encoding methyl-CpG-binding protein 2. Nat Genet, 1999. 23(2): p. 185–8.

3. Hagberg, B., et al., Rett syndrome: criteria for inclusion and exclusion. Brain Dev, 1985. 7(3): p. 372–3.

4. Demeter, K., Assessing the developmental level in Rett syndrome: an alternative approach? Eur Child Adolesc Psychiatry, 2000. 9(3): p. 227–33.

5. Percy, A.K., et al., Rett syndrome diagnostic criteria: lessons from the Natural History Study. Ann Neurol, 2010. 68(6): p. 951–5.

6. Berger-Sweeney, J., Cognitive deficits in Rett syndrome: what we know and what we need to know to treat them. Neurobiol Learn Mem, 2011. 96(4): p. 637–46.

7. Katz, D.M., et al., Rett Syndrome: Crossing the Threshold to Clinical Translation. Trends Neurosci, 2016. 39(2): p. 100–113.

8. Peters, S.U., et al., Distinguishing response to names in Rett and MECP2 Duplication syndrome: An ERP study of auditory social information processing. Brain Res, 2017. 1675: p. 71–77.

9. Foxe, J.J., et al., Automatic cortical representation of auditory pitch changes in Rett syndrome. J Neurodev Disord, 2016. 8(1): p. 34.

10. LeBlanc, J.J., et al., Visual evoked potentials detect cortical processing deficits in Rett syndrome. Ann Neurol, 2015. 78(5): p. 775–86.

11. Goffin, D., et al., Rett syndrome mutation MeCP2 T158A disrupts DNA binding, protein stability and ERP responses. Nat Neurosci, 2011. 15(2): p. 274–83.

12. Peters, S.U., R.L. Gordon, and A.P. Key, Induced gamma oscillations differentiate familiar and novel voices in children with MECP2 duplication and Rett syndromes. J Child Neurol, 2015. 30(2): p. 145–52.

13. Stauder, J.E., et al., The development of visual- and auditory processing in Rett syndrome: an ERP study. Brain Dev, 2006. 28(8): p. 487–94.

14. Stach, B.A., et al., Auditory evoked potentials in Rett syndrome. J Am Acad Audiol, 1994. 5(3): p. 226–30.

15. Mills, D.L., et al., Language experience and the organization of brain activity to phonetically similar words: ERP evidence from 14- and 20-month-olds. J Cogn Neurosci, 2004. 16(8): p. 1452–64.

16. Yoder, P.J., et al., Association between differentiated processing of syllables and comprehension of grammatical morphology in children with Down syndrome. Am J Ment Retard, 2006. 111(2): p. 138–52.

17. Riva, V., et al., Distinct ERP profiles for auditory processing in infants at-risk for autism and language impairment. Sci Rep, 2018. 8(1): p. 715.

18. Näätänen, R., Mismatch negativity: clinical research and possible applications. International Journal of Psychophysiology, 2003. 48(2): p. 179–188.

19. Ritter, W., et al., Memory reactivation or reinstatement and the mismatch negativity. Psychophysiology, 2002. 39(2): p. 158–65.

20. Young, E.D., Neural representation of spectral and temporal information in speech. Philos Trans R Soc Lond B Biol Sci, 2008. 363(1493): p. 923–45.

21. Peter, V., G. McArthur, and W.F. Thompson, Discrimination of stress in speech and music: a mismatch negativity (MMN) study. Psychophysiology, 2012. 49(12): p. 1590–600.

22. De Sanctis, P., et al., Auditory scene analysis: the interaction of stimulation rate and frequency separation on pre-attentive grouping. Eur J Neurosci, 2008. 27(5): p. 1271–6.

23. Schroger, E. and I. Winkler, Presentation rate and magnitude of stimulus deviance effects on human pre-attentive change detection. Neurosci Lett, 1995. 193(3): p. 185–8.

24. Kaufmann, W.E., et al., Social impairments in Rett syndrome: characteristics and relationship with clinical severity. J Intellect Disabil Res, 2012. 56(3): p. 233–47.

25. Delorme, A. and S. Makeig, EEGLAB: an open source toolbox for analysis of single-trial EEG dynamics including independent component analysis. J Neurosci Methods, 2004. 134(1): p. 9–21.

26. Molholm, S., et al., The neural circuitry of pre-attentive auditory change-detection: an fMRI study of pitch and duration mismatch negativity generators. Cereb Cortex, 2005. 15(5): p. 545–51.

27. De Sanctis, P., et al., Right Hemispheric Contributions to Fine Auditory Temporal Discriminations: High-Density Electrical Mapping of the Duration Mismatch Negativity (MMN). Front Integr Neurosci, 2009. 3: p. 5.

28. Butler, J.S., et al., Common or redundant neural circuits for duration processing across audition and touch. J Neurosci, 2011. 31(9): p. 3400–6.

29. Meng, X.-l., R. Rosenthal, and D.B. Rubin, Comparing correlated correlation coefficients. Psychological Bulletin, 1992. 111(1): p. 172–175.

30. Guthrie, D. and J.S. Buchwald, Significance testing of difference potentials. Psychophysiology, 1991. 28(2): p. 240–4.

31. Molholm, S., et al., Multisensory auditory-visual interactions during early sensory processing in humans: a high-density electrical mapping study. Brain Res Cogn Brain Res, 2002. 14(1): p. 115–28.

32. Näätänen, R., et al., The mismatch negativity (MMN) – A unique window to disturbed central auditory processing in ageing and different clinical conditions. Clinical Neurophysiology, 2012. 123(3): p. 424–458.

33. Bishop, D.V.M., M.J. Hardiman, and J.G. Barry, Is auditory discrimination mature by middle childhood? A study using time-frequency analysis of mismatch responses from 7 years to adulthood: Is auditory discrimination mature? Developmental Science, 2011. 14(2): p. 402–416.

34. Glass, E., S. Sachse, and W. von Suchodoletz, Development of auditory sensory memory from 2 to 6 years: an MMN study. Journal of Neural Transmission, 2008. 115(8): p. 1221–1229.

35. Cheour, M., et al., The auditory sensory memory trace decays rapidly in newborns. Scandinavian Journal of Psychology, 2002. 43(1): p. 33–39.

36. Glass, E., S. Sachse, and W. von Suchodoletz, Auditory sensory memory in 2-year-old children: an event-related potential study. NeuroReport, 2008. 19(5): p. 569–573.

37. Galbraith, G.C., M. Philippart, and L.M. Stephen, Brainstem frequency-following responses in Rett syndrome. Pediatric Neurology, 1996. 15(1): p. 26–31.

38. Bartha-Doering, L., et al., A systematic review of the mismatch negativity as an index for auditory sensory memory: From basic research to clinical and developmental perspectives: MMN as an index for auditory sensory memory. Psychophysiology, 2015. 52(9): p. 1115–1130.

39. Pekkonen, E., et al., Aging Effects on Auditory Processing: An Event-Related Potential Study. Experimental Aging Research, 1996. 22(2): p. 171–184.

40. Schroeder, M.M., W. Ritter, and H.G. Vaughan, The Mismatch Negativity to Novel Stimuli Reflects Cognitive Decline. Annals of the New York Academy of Sciences, 1995. 769(1 Structure and): p. 399–401.

41. Javitt, D.C., et al., Role of cortical N-methyl-D-aspartate receptors in auditory sensory memory and mismatch negativity generation: implications for schizophrenia. Proceedings of the National Academy of Sciences, 1996. 93(21): p. 11962–11967.

42. Rosburg, T. and I. Kreitschmann-Andermahr, The effects of ketamine on the mismatch negativity (MMN) in humans – A meta-analysis. Clinical Neurophysiology, 2016. 127(2): p. 1387–1394.

43. Blue, M.E., S. Naidu, and M.V. Johnston, Altered Development of Glutamate and GABA Receptors in the Basal Ganglia of Girls with Rett Syndrome. Experimental Neurology, 1999. 156(2): p. 345–352.

44. Blue, M.E., et al., Temporal and Regional Alterations in NMDA Receptor Expression in Mecp2-Null Mice. The Anatomical Record: Advances in Integrative Anatomy and Evolutionary Biology, 2011. 294(10): p. 1624–1634.

45. Magnusson, K.R., B.L. Brim, and S.R. Das, Selective Vulnerabilities of N-methyl-D-aspartate (NMDA) Receptors During Brain Aging. Front Aging Neurosci, 2010. 2: p. 11.

46. Pekkonen, E., et al., Auditory sensory memory impairment in Alzheimer's disease: an event-related potential study. Neuroreport, 1994. 5(18): p. 2537–40.

47. Gaeta, H., et al., The effect of perceptual grouping on the mismatch negativity. Psychophysiology, 2001. 38(2): p. 316–324.

48. Laptinskaya, D., et al., Auditory Memory Decay as Reflected by a New Mismatch Negativity Score Is Associated with Episodic Memory in Older Adults at Risk of Dementia. Frontiers in Aging Neuroscience, 2018. 10.

49. Lee, M., et al., A tale of two sites: Differential impairment of frequency and duration mismatch negativity across a primarily inpatient versus a primarily outpatient site in schizophrenia. Schizophrenia Research, 2018. 191: p. 10–17.

50. Magno, E., et al., Are auditory-evoked frequency and duration mismatch negativity deficits endophenotypic for schizophrenia? High-density electrical mapping in clinically unaffected first-degree relatives and first-episode and chronic schizophrenia. Biol Psychiatry, 2008. 64(5): p. 385–91.

51. Zhao, Y.T., et al., Loss of MeCP2 function is associated with distinct gene expression changes in the striatum. Neurobiol Dis, 2013. 59: p. 257–66.

52. Bishop, D.V., et al., Auditory development between 7 and 11 years: an event-related potential (ERP) study. PLoS One, 2011. 6(5): p. e18993.

53. Lipponen, A., et al., Auditory-evoked potentials to changes in sound duration in urethane-anaesthetized mice. Eur J Neurosci, 2019.

